# *Cacna1b* alternative splicing is linked to associative learning

**DOI:** 10.64898/2026.02.17.706441

**Authors:** Simrat Kaur Dhillon, Ava Cardarelli, Ashton Brennecke, Aaron Bradford, Alexandra Bunda, Forest MacKenzie, Vladimir Tkachev, Colin Call, Arturo Andrade

## Abstract

Voltage-gated Ca_V_2.2 channels are essential for neurotransmitter release throughout the nervous system including areas related to learning and memory like the hippocampus. Previous results have shown that Ca_V_2.2 channels are involved in cognitive processes. However, a link between alternative splicing of the *Cacna1b* (gene that encodes for Ca_V_2.2) pre-mRNA and cognitive processes has not been described. The *Cacna1b* pre-mRNA undergoes extensive cell-specific alternative splicing. In this body of work, we focus on the cassette exon 18a. Alternative splicing of exon 18a generates two splice variants, +18a-*Cacna1b* and Δ18a-*Cacna1b*. Exon 18a encodes a 21-amino acid sequence within the SYNaptic PRotein INTeraction (*synprint*) site. Splice variants containing exon 18a (+18a-Ca_V_2.2) show reduced cumulative inactivation and increased Ca²⁺ current density compared to splice variants lacking exon 18a (Δ18a-Ca_V_2.2), suggesting functional specialization. We previously showed that +18a-*Cacna1b* splice variants are enriched in cholecystokinin-expressing interneurons (CCK^+^INs). This neuronal type is strongly implicated in associative learning. Therefore, we tested whether alternative splicing of exon 18a contributes to associative learning. To test this hypothesis, we used genetically engineered mice that constitutively express either +18a-*Cacna1b* (+18a) or Δ18a-*Cacna1b* (Δ18a). We first validated that restricted splicing of exon 18a did not alter downstream alternative or constitutive spliced exons in the *Cacna1b* pre-mRNA, nor total Ca_V_2.2 protein levels. We then performed a comprehensive behavioral analysis that included assessment of associate learning. We found that in the trace fear conditioning task, +18a mice exhibited less freezing during the trace interval in both the acquisition and memory phases compared to WT mice. Whereas Δ18a mice showed enhanced freezing during the same intervals relative to WT mice. These bidirectional phenotypes reveal that exon 18a shapes aversive associative learning. Furthermore, exon 18a splicing did not influence spatial working memory, spatial navigation under stress, nociceptive responses in basal and inflammatory conditions, overall locomotion or exploratory behavior. These results suggest that the behavioral impact of exon 18a splicing is highly selective. Together, our findings identify alternative splicing of exon 18a as a molecular contributor to associative learning.

## INTRODUCTION

Ca_V_2.2 channels are critical for transmitter release in both the peripheral and central nervous system. (1) In the periphery, Ca_V_2.2 channels play a major role in the release of neurotransmitter from presynaptic terminals of nociceptors into the spinal cord, as well as, on the release of pro-nociceptive molecules from skin nerve endings. (2-4) In the central nervous system, Ca_V_2.2 channels dominate in dopamine transmission in the caudate putamen and nucleus accumbens. (5) Ca_V_2.2 channels are key for the release of neurotransmitter from cholecystokinin expressing interneurons (CCK^+^IN) located in the basolateral amygdala (6), and hippocampus (7-12). Cav2.2 channels, together with Ca_V_2.1 and Ca_V_2.3 channels, contribute to the release of glutamate from pyramidal neurons. (1) Because their ability to control transmitter release at key hippocampal synapses, a potential role of Ca_V_2.2 channels on learning and memory has been observed.

Previous studies using specific Ca_V_2.2 channel blockers and Ca_V_2.2 knock-out mice have suggested that this channel is important for cognitive processes. For example, intracerebroventricular infusion of the Ca_V_2.2 blocker, *ɷ*-conotoxin GVIA, resulted in deficits during novel object location and novel object recognition tasks, suggesting impairments in spatial and non-spatial short-term memory. (13) Furthermore, Ca_V_2.2 knock-out mice showed memory deficits in a social transmission of food preference task. (14) Similarly, Ca_V_2.2 knock-out mice exhibited impairments in spatial learning and memory in the Morris water maze test. (14) These results support that Ca_V_2.2 channel is linked to both spatial and non-spatial long-term memory. However, the cellular and molecular mechanisms that link Ca_V_2.2 channels to cognitive processes are poorly understood.

The gene that encodes the Ca_V_α_1_ pore-forming subunit for Ca_V_2.2, *Cacna1b*, is subject to cell-specific alternative splicing. This process is well-known to generate Ca_V_2.2 splice variants with distinct functions. We have previously characterized several sites of alternative splicing in the *Cacna1b* pre-mRNA: the cassette exons 18a, 24a, and 31a; and the mutually exclusive exons, 37a and 37b. (15) In previous studies, we found that 37a-Ca_V_2.2 splice variants are enriched in nociceptors, influence regulation of Ca_V_2.2 channels by μ-opioid receptors, and couple to the analgesic actions of spinal morphine. (16, 17) 37a-Ca_V_2.2 splice variants are also present in pyramidal neurons, control glutamate synaptic probability release and impact exploratory behavior. (18, 19) These studies provide proof of concept that cell-specific alternative splicing of *Cacna1b* impacts cell-specific functions and behavior.

In this body of work, we study the cassette exon 18a (e18a). Alternative splicing of these exon results in splice variants with (+18a-*Cacna1b*) and without (Δ18a-*Cacna1b*). This exon encodes a 21 amino acid sequence within the intracellular linker region between domain II and III (**Fig. 1A**). (20) Exon 18a is strongly conserved among mice, rats, and humans; and follows a strikingly similar pattern of expression among these species.(21) The e18a sequence falls within the SYNaptic PRotein INTeraction (synprint) region, a sequence that is relevant for the interaction of Ca_V_2.2 with proteins of the release machinery including the Soluble NSF Attachment Protein Receptor (SNARE) proteins. (12, 22) At the functional level, inclusion of e18a produces Ca_V_2.2 channels with reduced cumulative inactivation relative to those without e18a. (23) Furthermore, e18a-CaV2.2 splice variants channels show larger Ca^2+^ currents compared to Δe18a-Ca_V_2.2 splice variants in both heterologous expression systems and neurons. (21) All this combined evidence supports that e18a alternative splicing may play key roles in cells and tissues this splicing event occurs.

**Figure 1.**
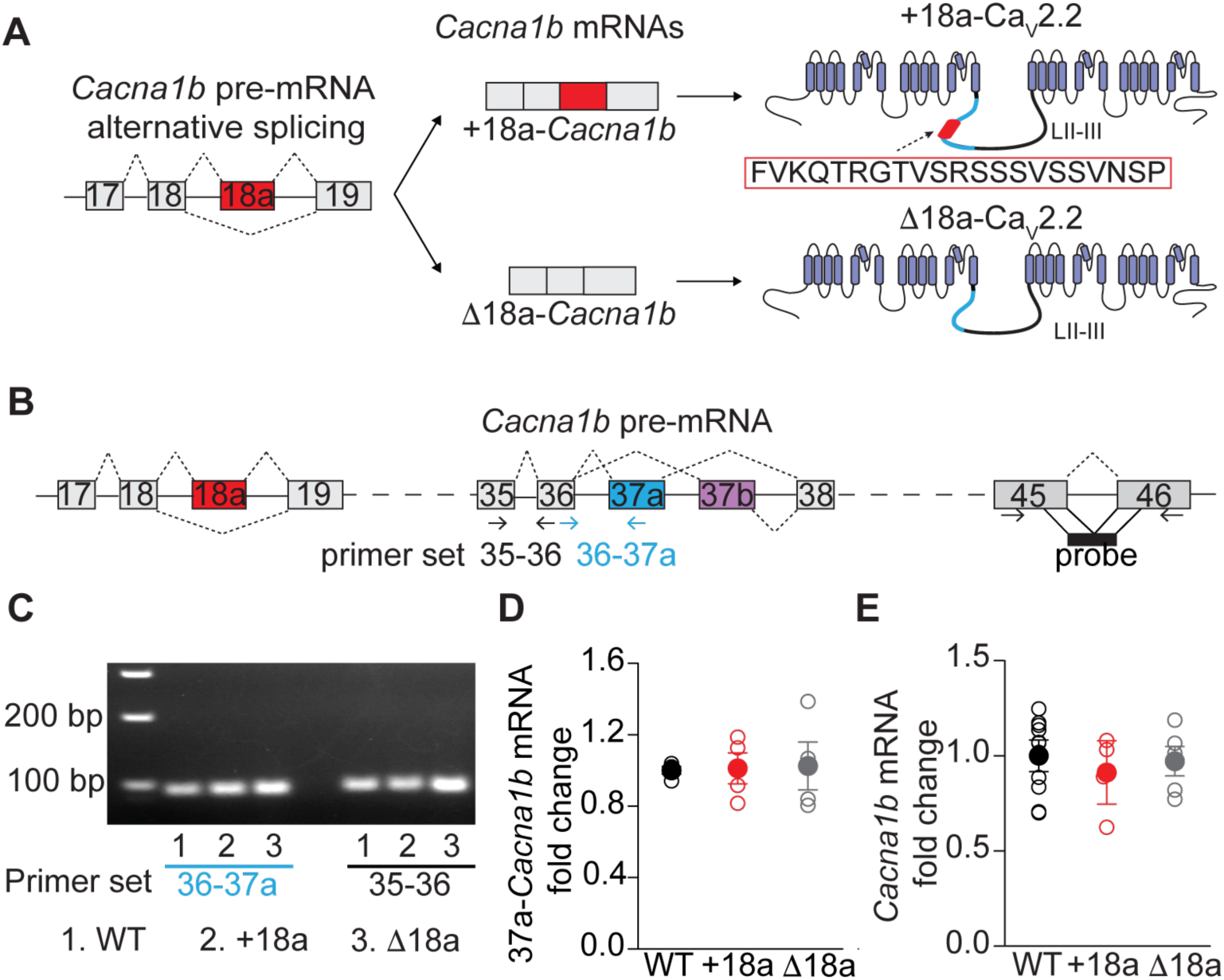
Restriction of splice choice at e18a preserves splicing and total Cacna1b mRNA levels. **A)** Schematic representation of alternative splicing of e18a in the *Cacna1b* pre-mRNA. Splicing of e18a results in two splice variants (+18a and Δ18a). Splice variants with +18a contain a 21 aminoacid sequence (red box) in the synprint site (approximate location highlighted in blue) in the linker between DII and DII (LII-III). **B**) Schematic representation of the *Cacna1b* pre-mRNA. Gray boxes represent constitutive exons and colored boxes alternative spliced exons. Continuous lines depict non-coding pre-mRNA sequences and long-dashed lines are sequences intentionally omitted. Short-dashed lines represent splice sites. Approximate location of primer sets used to amplify 35-36 and 36-37a splice junction are shown. Similarly, approximate location of a probe used to quantify total *Cacna1b* mRNA is shown. **C**) Representative gel electrophoresis of an RT-PCR used to amplify 35-36 and 36-37a fragments in whole-brain tissue from WT, +18a, and Δ18a adult mice. **D**) RT-qPCR for e37a in whole-brain tissue. Amplification for e37a was normalized to 35-36 amplifications for all conditions tested. Data are presented as mean and s.e.m. for fold change of e37a mRNA relative WT. Filled circles are the mean and empty circles represent values from whole brain tissue of individual mice. **E**) RT-qPCR for total *Cacna1b* mRNA in whole-brain tissue from WT, +18a, and Δ18a adult mice. Data are presented as mean and s.e.m. of fold change for *Cacna1b* mRNA relative to WT. Filled circles are mean and empty circles values from individual mice. *Cacna1b* mRNA levels were normalized to *Gapdh* mRNA.

We have previously shown that +18a-*Cacna1b* splice variants are enriched in CCK^+^INs in the hippocampus. (18) CCK^+^INs play critical roles in emotional processing and associative learning. (24-26) Therefore, we hypothesized that alternative splicing of e18a in the *Cacna1b* pre-mRNA is linked to associative learning. To test this hypothesis, we used previously generated mice with restricted 18a splicing. (21)These mice express either +18a-Ca_V_2.2 or Δ18-Ca_V_2.2 splice variants but not both (+18a and Δ18a, respectively) (21) First, we rule out that our genetic manipulation affected alternative and constitutive splicing of downstream exons and total protein levels for Ca_V_2.2. Second, we performed an extensive behavioral characterization to show that alternative splicing of e18a is linked to associative learning but not to other behaviors typically connected to Ca_V_2.2 function like nociception and locomotion. These results unveil a novel function of *Cacna1b* splice variants in learning and memory.

## METHODS

### Animals

#### Housing conditions and handling

All experimental procedures were performed in accordance with the Institutional Animal Care and Use Committees (IACUC) at the University of New Hampshire and Brown University. Male and female C57BL/6J mice (2–3 months old) of the following genotypes were used for all of our experiments: WT, +18a (*Cacna1b^tm3.1Dili^*) and Δ18a (*Cacna1b^tm4.1Dili^*) (21) Brain tissue from Ca_V_2.2-null mice (*Cacna1b^tm5.1Dili^*) served as negative controls for validating the specificity of anti-Ca_V_2.2 antibodies in Western-blot analysis (see Western Blot section) (2)Unless otherwise noted, mice were housed in same-sex groups under a 12 h light/dark cycle and were provided food and water ad libitum. Experimenters were blind to the genotype of all animals throughout testing and data analysis.

#### Δ18a and +18a mouse lines

The generation of the +18a and Δ18a mouse lines has been described previously. (21) Targeting constructs to constitutively include e18a (+18a allele) or eliminate e18a (Δ18a allele) were produced from the Sanger Bacterial Artificial Chromosome BMQ175b12. (27) To generate the +18a allele, the introns flanking e18a were removed; and e18a was fused to e18 and e19. To produce the Δ18a allele, a targeting construct was designed to delete the genomic region spanning from the intron immediately downstream of e18 through the intron immediately upstream of e19. Following homologous recombination in embryonic stem cells and blastocyst injection, chimeric founders were crossed to C57BL/6 mice to generate F1 heterozygotes. Homozygous progeny were backcrossed to C57BL/6 (Charles river) mice for at least six generations to ensure a congenic background. (21)

### RT-qPCR

Whole brains were rapidly dissected, flash-frozen in liquid nitrogen, and stored on dry ice until processing. Frozen tissue from WT, +18a, and Δ18a mice was pulverized under liquid nitrogen with a pre-chilled mortar and pestle. Approximately 30 mg of pulverized tissue was homogenized in RLT lysis buffer (Qiagen, 74134) and total RNA was extracted using Qiagen RNAeasy plus kit (Qiagen, 74134) according to manufacturer instructions. RNA was eluted in diethyl pyrocarbonate-treated water and 1 μg of total RNA was reverse-transcribed with oligo-dT primers using Superscript IV First-Strand Synthesis System (ThermoFisher Scientific, 18091050).

Relative abundance of e37a-containing *Cacna1b* transcripts was determined by RT-qPCR as described previously (18). Primers specific for the 36-37a splice junction (e36-37aF: 5’-CTGCGTGTTGCCGGATT; e36-37aR: 5’ACCTACGAGGGCAGTTCTT) were used in parallel with primers recognizing constitutive exons 35-36 (e35-36F: 5’ GGAAACATTGCCCTTGATGATG, e35-36R: 5’ CAGTGGCACTCCTGAACAATA), the latter serving as the internal references for normalization. Primer efficiency and specificity have been validated previously. (18) All RT-qPCR reactions were run on an ABI 7500 Fast Real-Time PCR system (Applied Biosystems) with the following conditions: 1 cycle 95 °C for 2 min, 45 cycles (95 °C for 15 s and 60 °C for 1 min). Each sample from at least five different mice per genotype (biological replicates) was run in triplicate (technical replicates). Ct values were determined by 7500 Software v2.3 (Applied Biosystems). Relative quantification of gene expression was performed with the 2-ΔΔCt method. Total *Cacna1b* mRNA quantification was performed with commercially available probes Mm01333678_m1 (ThermoFisher Scientific) targeting the last two exons using TaqMan® real-time PCR assays (ThermoFisher Scientific). Levels of mRNA were normalized to glyceraldehyde 3-phosphate dehydrogenase (*Gapdh*) using the probe Mm99999915_g1. Cycling conditions were similar to those described above.

### Western blot

Whole brains of 2-3 month mice were rapidly dissected, flash frozen in liquid nitrogen and homogenized in ice-cold 1x RIPA buffer (50 mM Tris HCl, pH 7.5; 150 mM NaCl; 1% Igepal CA-630; 0.5% sodium deoxycholate; and 0.1% sodium dodecyl sulfate) supplemented with one cOmplete^TM^ EDTA-free protease-inhibitor tablet per 10 mL (Millipore-Sigma, 11873580001). Homogenates were incubated on ice for 30 minutes, re-homogenized and centrifuged at 1,000 x g for 10 min at 4°C to remove cell debris. The supernatant was then centrifuged 12,000 x g for 20 min at 4°C; the resulting lysate was reserved for analysis. Protein concentrations were determined with the bicinchoninic acid (BCA) assay (ThermoFisher, 23227). 20 μg of sample was loaded in electrophoresis gel. For SDS-PAGE, samples were mixed 1:1 with 2x reducing sample buffer (100 mM Tris-HCl, pH 6.8; 4% SDS; 0.2 % bromophenol blue; 20% glycerol; 200 mM dithiothreitol) and denatured for 20 minutes at room temperature.

Proteins were resolved using a 6% polyacrylamide gel and a 4 % stacking gel. Electrophoresis running buffer (25 mM Tris, 192 mM glycine, 0.1 % SDS). WesternSure® Pre-stained Chemiluminescent Protein Ladder (Li-Cor, # 926-98000) and samples were loaded into the BIO-RAD mini-PROTEAN 3 system (Bio-Rad). Samples were run at 10 mA for 90 minutes, followed by 20 mA for another 30 minutes. Proteins were transferred to a polyvinylidene difluoride (PVDF) membrane using ice-cooled transfer buffer (25 mM Tris, 192 mM glycine, 0.1 % SDS, 20 % methanol). Prior to transfer, the PVDF membrane and gel were equilibrated for 20 minutes in 100% methanol and 1X transfer buffer, respectively, for 20 minutes. Transfer was performed at 100 V for 1 h on ice. Following transfer, the membrane was rinsed with 1 % PBS-Tween (PBS-T) and air-dried overnight. The PVDF membrane was reactivated in 100% methanol for one minute followed by a brief PBS-T rinse step. The membrane was incubated with REVERT total protein stain (Li-Cor #926-11014) for 5 minutes. Membranes were rinsed twice with wash solution for 30 seconds followed by a brief PBS-T rinse and subsequent storage in light-protected PBS-T. Membranes were imaged using an Odyssey CLx imaging system on the 700 nm channel. Next, membrane was blocked for 1 h with 5% BSA in PBS-T at RT and washed three times for 3 min in PBS-T. Anti-Ca_v_2.2 primary antibody (Alomone Rabbit anti-Ca_v_2.2, #ACC-002) was prepared 1:100 with 1% BSA in PBS-T and 15 μL 10 % NaN_3_ for a final concentration of 0.425 µg/mL. Primary antibody incubation was carried out for 1 hour at RT, followed by three washes in PBS-T. The peroxidase-conjugated F(ab)2 fragment donkey Anti-Rabbit IgG (H+L) (Jackson Immunoresearch, 168969) was prepared 1:15,000 in 1% BSA with PBS-T and incubated for 1 hour at RT. Membranes were washed 3 times for 10 min. Total protein stain and chemiluminescence images were analyzed using ImageJ software (NIH http://imagej.nih.gov/ij, v.1.51j8). Images were converted to 8-bit grayscale for analysis. To compare Ca_V_2.2 levels in WT, +18a, and Δ18a mice, chemiluminescence images were normalized to total stain fluorescence.

### Behavioral assays

#### Hargreaves Plantar Test

Mice were placed individually in clear acrylic chambers (IITC Life Science, 435) positioned on an elevated transparent glass platform. Mice were allowed to habituate to the behavioral testing room and scorer for 1 h. A focused radiant-light source (IITC Life Science, 390) was calibrated to result in the average mouse reacting after 10 s of exposure. During habituation, the light source was periodically moved around to allow the mice to familiarize with its presence. After habituation, the light source was aimed at the plantar surface of the left rear paw. A cut off time was set up at 20 s, and an inter-trial period of at least 5 minutes was given to each mouse to prevent desensitization. 3-5 successful trials per mouse was performed to measure the latency to withdrawal. For experiments involving Complete Freund’s adjuvant (CFA) mice were deeply anesthetized using 2-3% of isoflurane and 25 μL of CFA was injected in the left paw. For capsaicin experiments, mice were deeply anesthetized with isoflurane and lightly restrained, 0.1% w/v capsaicin was injected in the left paw.

#### Open Field Maze Test

Mice were habituated to the testing room for 30 minutes prior to behavioral assay. Each animal was placed in a square open-field arena (45 x 45 cm; opaque Plexiglass, 40 cm high) and allowed to freely explore for 5 min under uniform ambient lighting (20 lux). Mice were filmed using a video camara and tracked with EthoVision XT 11.5 software (Noldus Information Technologies, Inc). Total distance traveled, frequency of entry into the inner zone, and percent of time spent in the inner zone were calculated using EthoVision XT.

#### Elevated Plus Maze Test

Mice were placed for 10 minutes on an elevated-plus maze (EPM) consisting of four arms of 65 cm length, two of these arms were walled (closed arms) and two remained open (Harvard Apparatus, 76-0075). After each trial, mice were placed back in their home cages. Mice were video recorded and tracked with EthoVision XT 11.5. Total distance traveled, frequency of entries into open arms, and the percent of time spent in the open arms of the EPM was calculated with Ethovision XT.

#### Barnes Maze Test

The Barnes maze consisted of circular platform (92 cm diameter) elevated 90 cm above the floor containing 20 equidistant 5-cm-diameter holes around its perimeter; only one of these holes (target hole) was connected to an escape box affixed underneath the platform. The maze was brightly lit from the top to induce mild stress (1300 lux) and covered with white curtains. Visual cues of distinct colors and shapes were placed around the maze to guide mice to find the target hole. At the beginning of each trial, mice were placed in the center of the apparatus with the aid of an opaque cylinder. After 8 s acclimation period, the cylinder was removed, and mice were allowed to explore for up to 5 min to locate and enter the target hole. If the target was not found within 5 min, the experimenter gently guided the mouse to the hole and allowed it to remain in the escape box for 2 min before returning it to the home cage. Each animal completed two trials separated by a thirty-minute interval for seven consecutive days.

Trials were recorded with and scored for the total distance traveled, average velocity, latency to target hole, cumulative duration in target zone, and number of errors EthoVision XT 11.5. Search strategies were scored manually by the experimenter; the strategy employed by the animal during each trial was classified as either direct, serial, or random. Trials were classified as “direct” when the animal checked only one or two holes adjacent to the target hole before locating the target hole, “serial” when the animal searched each hole in a sequential manner along the perimeter of the apparatus before locating the target hole, and “random” when the strategy employed followed no obvious pattern. Trials were also classified as “random” when the animal failed to locate the target hole within the designated 5-minute trial period.

#### Y-maze Test

The Y-Maze apparatus consists of a Y-shaped maze with three arms evenly spaced at 120°. At the start of each trial, the test animal is placed in the center of the Y-maze and allowed to freely explore the three arms for 10 min under dim light. Mice were video recorded and tracked with EthoVision XT 11.5 This test assesses spatial working memory via percent alternations. This is calculated as the percent of spontaneous alternations, defined as successive entries into all three arms without revisiting any arm (e.g. A, B, C), relative to the maximum number of possible of alternations, defined as the total arm entries minus 2.

#### Trace Fear-Conditioning

During day I (acquisition phase), test animals were placed in their respective chambers and allowed to freely explore for 120 s before undergoing five conditioning trials. Each trial consisted of a 20 s tone (2.9 kHz, 80 dB), a 20 s trace interval, and then a 2 s 0.8 mA foot shock followed by an inter-trial interval (ITI) of 120 s. During day II (contextual phase), the test animals were returned to the same context as day I and observed for 300 seconds without the presentation of either the tone or shock. During day III (tone and trace memory phase), the animals underwent the same testing protocol as in day I in a novel context, but without foot shock presentation. Freezing behavior (complete immobility except for respiration) was measured during each phase using AnyMaze video tracking software (Maze Engineers).

### Statistical analysis

To assess whether our data showed normal distribution, we performed the Shappiro-Wilk test for normality. T-test or ANOVA analysis was performed when Shappiro test showed p > 0.05 (normal distribution). Tukey-Honest Significant Difference (Tukey-HSD) was performed as post-hoc test for ANOVA. For not normally distributed samples, Kruskal-Wallis test was used to compare the medians. As indicated throughout the text, generalized linear mixed model (GLMM) was used to analyze the effects of our units of analysis on a given outcome. To ensure the appropriateness of our GLMMs, we performed convergence analysis and residual diagnostic for hierarchical regression models using the R package DHARMa. All our models showed convergence equal to 0 and no significant problems were detected using DHARMa, suggesting that our models appropriately explained the observed variance. All statistical analysis was performed using the R packages rstatix, glmmTMB, emmeans, afex. Search strategy in the Barnes Maze was compared using Bayesian logistic regression (BRM) models with the R package brm.

## RESULTS

### Constitutive and alternative splicing of the Cacna1b pre-mRNA remain intact in +18a and Δ18a mice

To determine the functional role of the alternatively spliced e18a in the *Cacna1b* pre-mRNA, we used two mouse lines that expressed either *Cacna1b* with e18a (+18a) or without (Δ18a). These mouse lines were created using homologous recombination. (18, 21) To rule out off-target effects derived from manipulating 18a splicing in the *Cacna1b* pre-mRNA, we performed an extensive validation of our mouse genetic models. We quantified the alternative spliced exon 37a and the constitutive exons 45-46, both located downstream of 18a, in whole brains of WT, +18a and Δ18a mice (**Fig. 1B**). To quantify 37a, we used two sets of primers, 35-36 and 36-37a, that have previously been validated for specificity and efficiency (**Fig. 1B-C** and (18)). Both sets of primers amplified the expected products in brain samples from the three genotypes (**Fig. 1C**). Amplicons of 35-36 primers were used for normalization. We found that the levels of 37a-*Cacna1b* mRNA is similar among WT, +18a and Δ18a mice (ANOVA, F_2,9_ = 0.017, p = 0.98. **Fig. 1D**). Taken together, our results suggest that genomic strategy to restrict e18a alternative splicing does not affect usage of other alternatively spliced exons in the *Cacna1b* pre-mRNA.

We next compared constitutive splicing of *Cacna1b* pre-mRNA in WT, +18a and Δ18a mice using RT-qPCR with a probe directed to the constitutive exon junction 45-46 (**Fig. 1B**). This qPCR was normalized to the constitutive gene *Gapdh*. No differences were found in the amount of e45-46 among the three genotypes (F_2,20_ = 2.079, ANOVA, p = 0.154. **Fig 1D**). Because 45-46 are spliced in all *Cacna1b* mRNAs, we can also conclude that the total *Cacna1b* mRNA levels are similar among WT, +18a and Δ18 mice. These results suggest that the genomic manipulation to restrict e18a splicing does not disrupt constitutive splicing of other exons and total *Cacna1b* mRNA in whole brain.

### Protein levels for Ca_V_2.2 are unaffected in +18a and Δ18 mice

Ca_V_2.2 protein levels were quantified in whole-brain samples of WT, +18a and Δ18a mice using semi-quantitative western blot (WB) (**Fig. 2**). The specificity of anti-Ca_V_2.2 antibodies was validated by comparing samples from WT and Ca_V_2.2-null mice in WB. As expected, we observed a band of ∼250 KDa in WT mice, but not Ca_V_2.2-null mice (**Fig. 2A**). Total stain protein was used as loading control. (28)We found no differences in the protein levels for Ca_V_2.2 channels in brains of WT, +18a and Δ18a mice (F_2,29_ = 0.424, ANOVA, p = 0.666). No interaction was found between sex and genotype (F_2,29_ = 0.23, ANOVA, p = 0.85) **Fig. 2B** and **2C**). Our results strongly suggest that our genomic strategy does not result in changes in total Ca_V_2.2 protein levels. Thus, our mouse models are appropriate to assess the function of +18a-Ca_V_2.2 and Δ18a-Ca_V_2.2 splice variants at the behavioral level.

**Figure 2.**
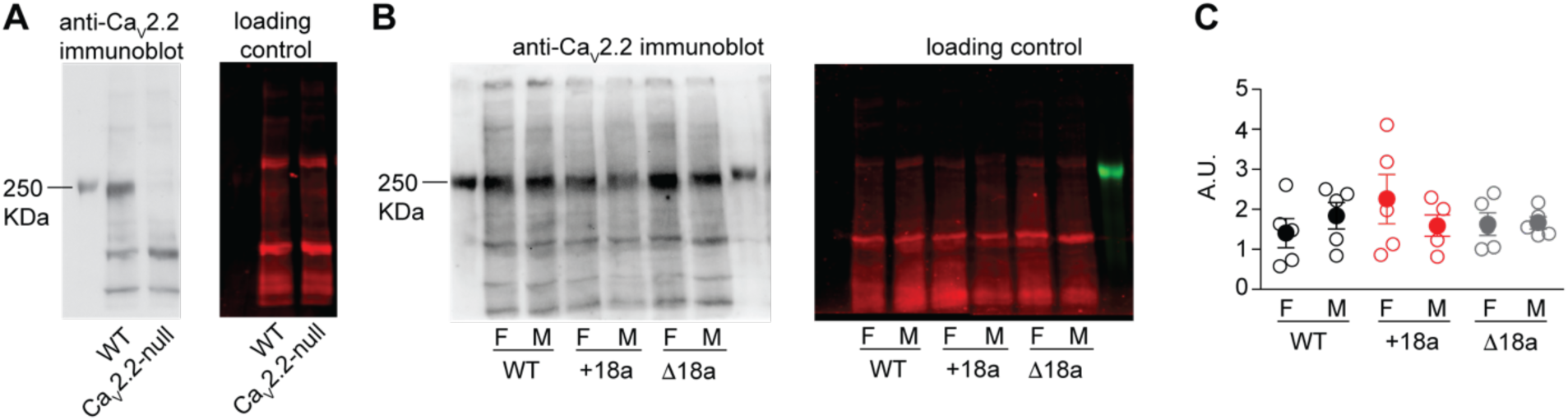
Disruption of 18a splicing did not change total Ca_V_2.2 protein levels in males and females. A) Anti-Ca_V_2.2 antibody validation. **B** and **C**) Ca_V_2.2 protein levels in brain of WT, +18a- and Δ18a mice. C) Mean (filled symbols) ± s.e.m and data from individual mice (empty symbols). F = females, M = males.

### Cacna1b alternative splicing of e18a is linked to aversive associative learning but not to working memory or spatial navigation under stress

Ca_V_2.2 channels, together with Ca_V_2.1 and Ca_V_2.3, control the release of glutamate from excitatory synapses onto pyramidal neurons (29, 30), whereas Ca_V_2.2 channels are the major presynaptic calcium channel controlling the release of GABA from CCK^+^INs in the hippocampus. (7-9, 11, 12) Our previous studies have shown that +18a-*Cacna1b* splice variants are enriched in CCK^+^INs and Δ18a-*Cacna1b* splice variants are dominant in pyramidal neurons in the hippocampus. (18) CCK^+^INs are key players in associative learning. (24-26) Given the pattern of expression for +18a and Δ18a-*Cacna1b* splice variants, we tested if there is a link between these splice variants and aversive associative learning using the trace fear conditioning (TFC) task. In TFC, a neutral conditioned stimulus (tone) is followed-after a stimulus-free temporal gap (20 s trace)- by an unconditioned stimulus (foot shock) (**Fig. 3A**). Successful acquisition of associative learning is quantified as the amount of freezing during the tone or the trace period. Our TFC paradigm was carried out across three days (see methods section). Day I (acquisition phase, **Fig. 3A**), day II (context, **Fig. 4**), and day III (tone and trace memory phase, **Fig. 5A**). We measured the percentage of freezing during baseline, tone presentation, trace period, and intertrial interval for days I and III. For day 2, freezing to the context was measured for 5 min.

**Figure 3.**
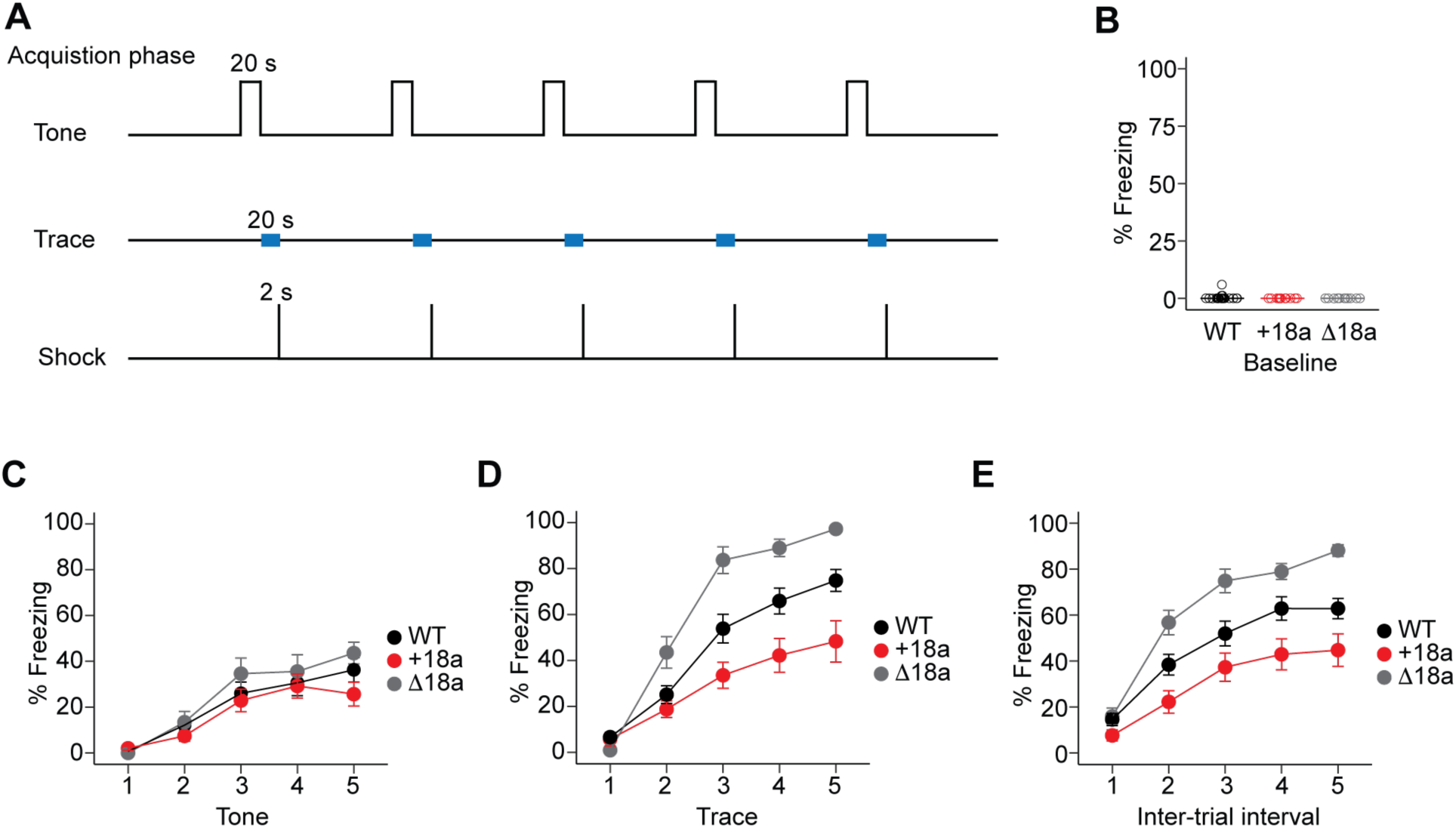
Alternative splicing of e18a impacts freezing during the trace and inter-trial interval periods in the acquisition phase of the trace fear conditioning task. **A**) Schematic representation of the protocol used in the acquisition phase of the trace fear conditioning task. After a 120 s baseline, five 20 s tones were delivered with a 20 s trace period followed by 2 s of a 0.8 mA shock. **B**) Freezing measured during the baseline period. Data represent mean (filled symbols) ± s.e.m and values from individual mice (empty symbols). **C**) Freezing during the five-tone presentation mean ± s.e.m are presented for the three genotypes. **D**) Freezing during the trace periods between tone and shock presentation, mean ± s.e.m. **E**) Freezing during the intertrial interval after shock presentation and before tone delivery of the next trial, mean ± s.e.m.

**Figure 4.**
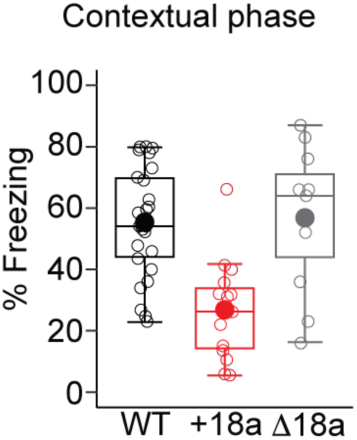
Constitutive inclusion of e18a impairs contextual fear conditioning. Freezing to the context was measured 24 hrs after the acquisition phase for mice of the three genotypes. Data are shown as mean (filled symbols), individual mice (empty symbols), interquartile range (box), median (horizontal line), and upper and lower whiskers, and outliers.

**Figure 5.**
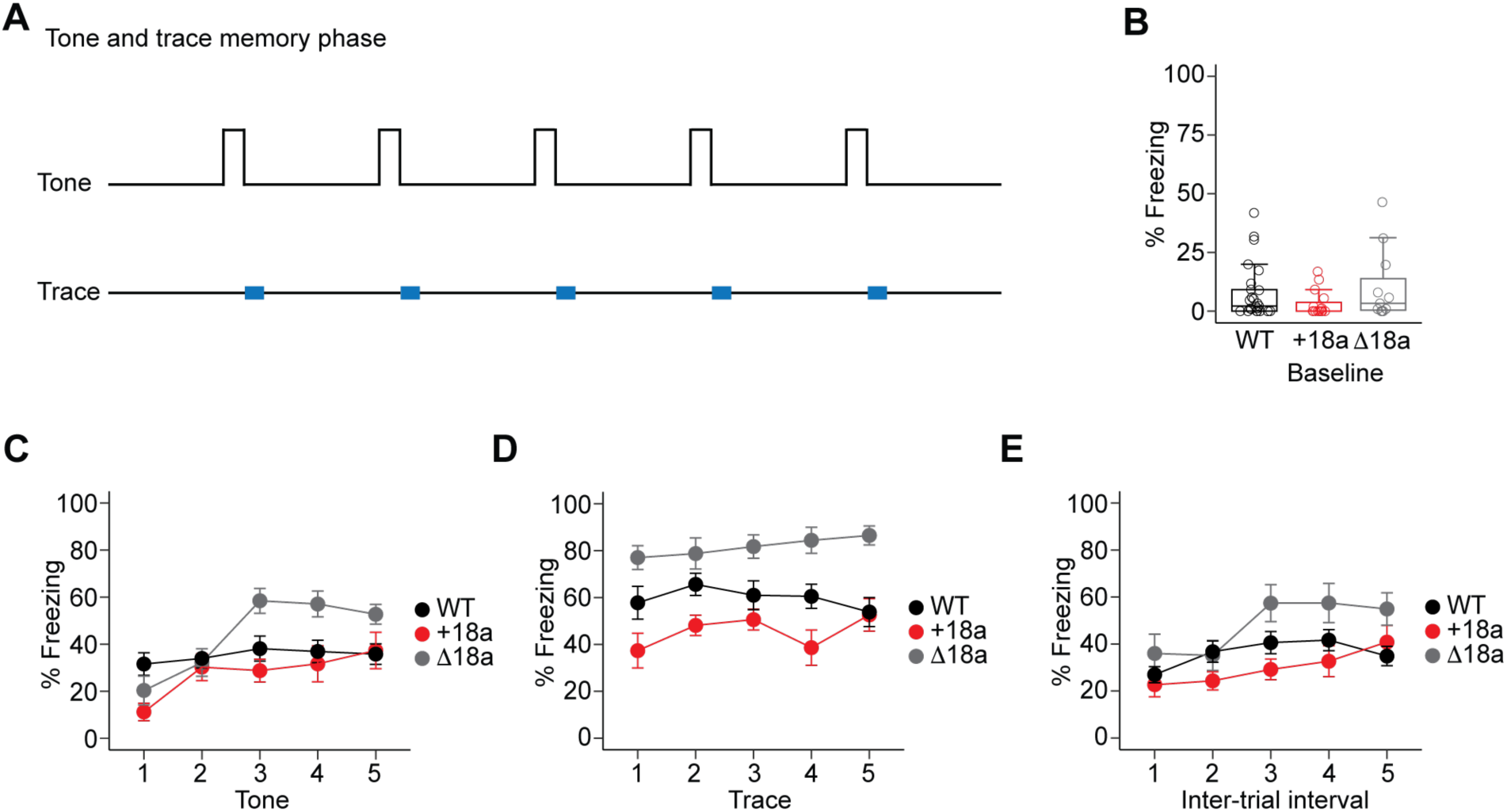
Alternative splicing of e18a impacts freezing during the trace period in the tone and memory phase of the trace fear conditioning task. **A**) Schematic representation of the protocol used in the trace and memory phase of a trace fear conditioning task. After a 120 s baseline, only five 20 s tones were delivered. The trace period was defined as the 20 s following the tone (blue rectangles). **B**) Freezing measured during the baseline period. Data are shown as mean (filled symbols), individual mice (empty symbols), interquartile range (box), median (horizontal line), and upper and lower whiskers, and outliers. **C**) Freezing during the five-tone presentation mean ± s.e.m are shown for the three genotypes. **D**) Freezing during the trace periods between tone and shock presentation, mean ± s.e.m. **E**) Freezing during the inter-trial interval after shock presentation and before tone delivery of the next trial, mean ± s.e.m.

During baseline in day I, we observed that only 5% of animals showed < 5% freezing suggesting that none of the environmental conditions in the chamber and testing room induce freezing (**Fig. 3B**). During the tone presentation and as expected, we observed that mice from the three genotypes showed a linear increase in freezing to the tone (β = 2.01, SE = 0.20, z = 10.02, p < 0.0001. GLMM. **Fig. 3C**). However, WT mice freeze similarly to +18a mice (β = -0.14, SE = 0.26, z = -0.53, p = 0.60. GLMM. **Fig. 3C**) and to Δ18a mice (β = 0.26, SE = 0.29, z = 0.96, p = 0.36. GLMM. **Fig. 3C**). These results suggest that the basic association between the tone and shock is intact in mice from the three genotypes. Interestingly, during the trace period we observed that +18a mice froze less compared to WT (β = -0.87, SE = 0.31, z = -2.78, p = 0.005. GLMM. **Fig. 3D**), by contrast Δ18a mice froze significantly more than WT (β = 0.92, SE = 0.35, z = 2.61, p = 0.009. GLMM. **Fig. 3D**). We also observed significant differences on how mice froze across the trace periods (genotype x trace period interaction). We observed that +18a mice exhibited a more gradual increase in freezing behavior across trace periods compared to WT (β = -1.12, SE = 0.31, z = -3.55, p = 0.0004. GLMM. **Fig. 3D**), whereas Δ18a showed a steeper increase in freezing behavior across trace periods compared to WT mice (β = 1.20, SE = 0.37, z = 3.26, p = 0.001. GLMM. **Fig. 3D**). These results suggest that +18a mice have defects in developing a temporal expectation to threat compared to WT mice, whereas Δ18a mice have an enhanced ability to encode the timing between the tone and shock. Consistent with the trace period, we observed significant differences during the inter-trial interval. WT mice froze more than +18a mice (β = -0.84, SE = 0.34, z = -2.48, p = 0.013. GLMM), but less than Δ18 mice (β = 0.90, SE = 0.38, z = 2.39, p = 0.017. GLMM. **Fig. 3E**). Taken together, these results show that *Cacna1*b alternative splicing in e18a influences the temporal association between the tone and shock during the acquisition phase of TFC.

We next assessed contextual fear conditioning (day II) by measuring freezing of mice from the three genotypes in the same context of the acquisition phase, but without shock or cue. We measured percent of freezing for a period of 5 min 24 h after the acquisition phase. We found a main effect of genotype in the percent of freezing (F_2,49_ = 13, p < 0.00003. One-way ANOVA. **Fig. 4**). Post-hoc analysis with Tukey HSD revealed that +18a mice froze significantly less than WT mice (p = 0.00005. **Fig. 4**) and Δ18a mice (p = 0.0005. **Fig. 4**). No differences were detected between WT and Δ18a mice (p = 0.97). These results suggest that +18a mice form weaker contextual associations compared to WT and Δ18a mice further supporting the role of *Cacna1b* alternative splicing on associative learning.

To assess for tone and trace memory (day III), we measured freezing 48 h after the acquisition phase in a different context but only the tone was delivered without the shock (**Fig. 5A**). Baseline measurements show that freezing behavior was not different among mice from three genotypes (% median freezing, IQR: WT = 2.17, 9.17, n = 25; +18a = 0, 5.5, n = 15, Δ18a = 3.33, 13.4; χ² = 3.30, df = 2, p = 0.19, Kruskal-Wallis test. **Fig 5B**). No statistically significant differences were observed in the total amount of freezing across tones between WT and +18a mice (β = -0.29, SE = 0.26, z = -1.12, p = 0.26, GLMM. **Fig. 5C**) or between WT and Δ18a (β = 0.44, SE = 0.28, z = 1.57, p = 0.11, GLMM. **Fig. 5C**). However, post-hoc analysis using estimated marginal means showed that Δ18a mice froze significantly more compared to WT and +18a mice during tones 3, 4 and 5 (all p < 0.05, Tukey’s HSD. **Fig. 5C**). During the trace period, WT mice froze significantly less compared to Δ18a mice (β = 1.07, SE = 0.31, z = 3.45, p = 0.00055. GLMM. **Fig. 5D**). Our analysis showed a trend for WT mice to freeze more compared to +18a mice (β = -0.52, SE = 0.28, z = -1.90, p = 0.056, GLMM. **Fig. 5D**). Interestingly, no significant differences were found during the intertrial interval among the three genotypes (all p > 0.05, GLMM. **Fig 5E**). These results suggest that *Cacna1b* alternative splicing is linked to the formation of associative memory with a temporal component.

Previous results have shown that pharmacological block of Ca_V_2.2 with ω-conotoxin GVIA in hippocampus impairs spatial short-term memory (13), but a potential role for splice variants +18a-*Cacna1b* and Δ18a-*Cacna1b* has not been tested. To assess a potential link between these splice variants and spatial short-term memory, we measured the performance of mice from the three genotypes in the Y-maze test. The Y-maze apparatus consists of three arms at an angle of 120°. In this test, we assess the mouse willingness to explore new environments by measuring spontaneous alternations among the three arms of the Y-maze. Unique alternations (triads of visits to the arms) correlate with short-term spatial memory. In our studies, we found no statistically significant difference in the percentage of alternations among the three genotypes (% alternations ± SEM: WT = 61.1 ± 0.96, n = 45; e18a = 60.4 ± 1.35, n = 33, D18a = 58.9 ± 1.25; F2,94 = 0.71, p = 0.49, One-way ANOVA. **Fig. 6A**). To rule out potential defects in the ability of the mice to visit the three arms, we also measured distance traveled throughout the Y-maze. We found no statistically significant difference in distance traveled among the three genotypes (Total distance traveled in m ± SEM: WT = 34.4 ± 1.28, n = 45; e18a = 34.1 ± 1.33, n = 33, Δ18a = 34.9 ± 1.52; χ² = 0.13, df = 2, p = 0.93, Kruskal-Wallis test. **Fig. 6B**). These results suggest that alternative splicing of e18a is not linked to short-term spatial memory and further confirms that overall locomotion is preserved among the mice from the three genotypes.

**Figure 6.**
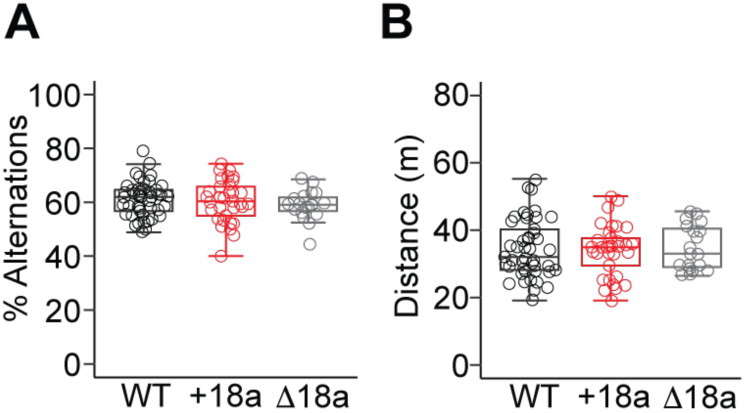
*Cacna1b* alternative splicing of exon 18a does not impact short-term spatial learning and memory. **A**) Comparison of per cent of spontaneous alternations in the Y-maze among mice from the three genotypes. Data are shown as mean (filled symbols), individual mice (empty symbols), interquartile range (box), median (horizontal line), and upper and lower whiskers, and outliers. **B**) Comparison of distance traveled in the Y-maze among mice from the three genotypes. Data are shown as mean (filled symbols), individual mice (empty symbols), interquartile range (box), median (horizontal line), and upper and lower whiskers, and outliers.

Behavioral studies using the Morris Water Maze (MWM) test and social transmission of food preference showed that Ca_V_2.2-knock out mice have impaired spatial learning and memory (14). To assess if alternative splicing of e18a cassette exon impacts spatial learning and memory under stress, we used an assay similar to the MWM test, the Barnes Maze test. In this test, the apparatus consists of an elevated circular platform with 20 holes and only one contains an escape box (31). The platform is brightly illuminated from the top, which provides a mild stressful stimulus for the mice. Around the platform, several visual cues are placed. Mice use these visual cues to find the hole containing an escape box (target) using their spatial navigation. Mice were trained on the maze for 7 days. We measured several parameters to assess learning and memory including latency to approach the hole with the escape box, distance traveled, the total number of errors, and the search strategy. For latency to approach the hole with escape box, we found that mice regardless of the genotype show a significant linear decrease in latency over time (β = -0.89, SE = 0.11, z = –7.76, p = 8.8 x 10^-15^, GLMM. **Fig. 7A**), consistent with learning. However, no significant effect of genotype or genotype x day interaction was observed (all p values > 0.05. **Fig. 7A**), suggesting similar learning trajectories among the three genotypes. These observations were confirmed with distance traveled, mice showed a significant linear decrease in distance travel over time regardless of the genotype (β = –0.94, SE = 0.1, z = -9.01, p = 2 x 10-^16^, GLMM. **Fig. 7B**), but no significant differences were associated with genotype or genotype x day interaction (all p values > 0.05. **Fig. 7B**). To analyze in more detail memory performance of mice from the three genotypes in the Barnes Maze, we compared the total number of errors over time defined as the sum of target revisits, non-target first visits, and non-target revisits. We found a significant linear decrease of total errors over time in mice from the three genotypes (β = –0.80, SE = 0.11, z = -7.47, p = 8.25 x 10^-14^, GLMM. **Fig. 7C**), but no significant effects of genotype or genotype x day interaction on total errors (all p values > 0.05. **Fig. 7C**). These results suggest that *Cacna1b* alternative splicing of e18a does not impact spatial navigation in the Barnes Maze test, and thereby spatial learning and memory.

**Figure 7.**
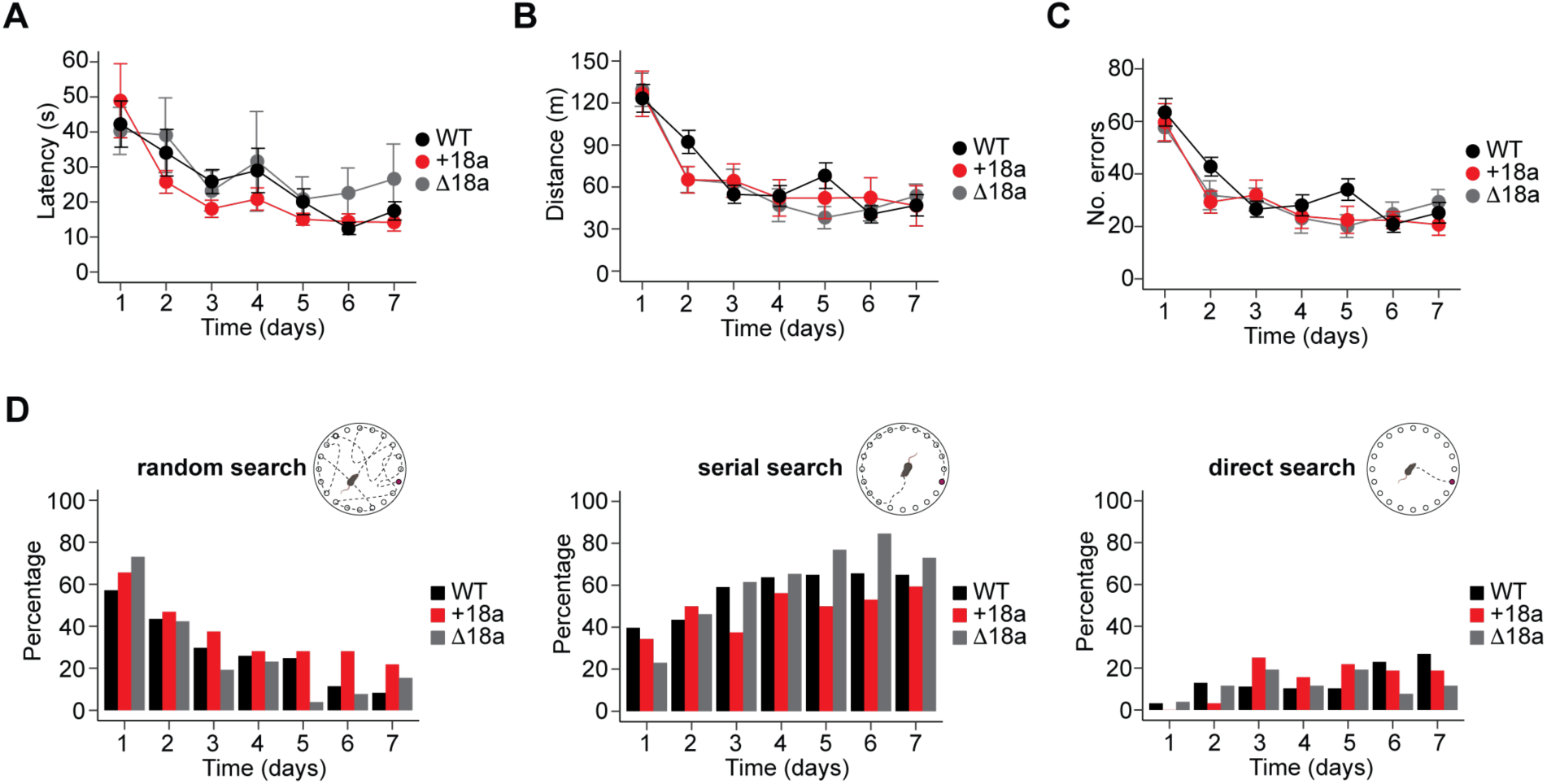
*Cacna1b* alternative splicing of exon 18a does not impact spatial navigation under mild stressful conditions. **A)** Latency to approach the escape box across seven days. **B**) Distance traveled (efficiency path) across seven days of training. **C**) Number of total errors (target revisits, non-target first visits, and non-target revisits) across seven days. **A**, **B**, and **C** are presented as mean and s.e.m. **D**) Types of search strategy (random, serial or direct, *inset*) that mice implemented to identify the escape box in the Barnes Maze. Data are presented as the percent of mice that predominantly exhibited either random, serial or direct search strategy across days. Note the mice from the three genotypes shifted from random to either serial or direct as training days increased.

For completion, we also analyzed the search strategy that mice use to find the target hole in the BM. Search strategy was categorized into three different types, random, serial and direct. We used a “winner takes it all” approach, where the dominant search strategy per trial was assigned to each mouse. To assess the likelihood that mice switched from random to a non-random search strategy (serial or direct) across day and genotypes, we fit a Bayesian logistic regression (BRM) model. As expected, and regardless of the genotype, we found a clear increase in the probability of non-random search strategy as the number of days increased. Starting from day 3, the main effects of day were positive, with day 3 showing an estimated log-odds of 1.64 (95% CI: [0.72, 2.57], BRM, **Fig. 7D**), increasing further on day 7 to 3.73 (95% CI: [2.54, 5.05], BRM, **Fig. 7D**). We did not find main effects from genotype on the probability of non-random search across days. These results suggest that mice from the three genotypes implement similar search strategies to find the hole with the escape box in the Barns Maze, indicating that they show similar learning. These results support the lack of link between *Cacna1b* alternative splicing in e18a and spatial navigation under mildly stressful conditions.

### Alternative splicing of exon 18a is dispensable for nociceptive behavior during inflammation

Previous studies have extensively shown that Ca_V_2.2 channels are critical to mediate nociceptive responses to thermal stimuli in basal conditions and after inflammation (4, 32, 33). Changes in alternative splicing for the *Cacna1b* pre-mRNA have been reported in neuropathic pain and inflammatory pain models (34, 35). *Cacna1b* splice variants of the mutually exclusive exons 37a and 37b have been linked to nociceptive behavior (34) and opioid peripheral analgesia (16, 17). However, the role of splice variants with or without e18a in nociception has not yet been investigated. To address this, we assessed thermal nociception using the Hargreaves test as previously described (16). 24 h after this initial testing, animals were injected with Complete Freund’s Adjuvant (CFA) to induce inflammation in one paw. The next day, animals were tested for thermal nociception every other day for 6 days in both paws (**Figure 8**). First, we compared withdrawal thresholds among WT, +18a, and Δ18a mice from both paws. A small but significant effect of genotype on thermal threshold was found (two-way ANOVA, F_2,90_ = 10.04, p = 0.046). No effect of paw on thermal threshold was observed (two-way ANOVA, F_1,90_ = 0.009, p = 0.92) or interaction of paw by genotype (two-way ANOVA, F_2,90_ = 0.403, p = 0.67). However, post-hoc comparisons using Tukey’s HSD test did not identify any statistically significant differences among the three genotypes (*all p* > 0.05. **Fig. 8A**). These results show that +18a-*Cacna1b* and Δ18a-*Cacna1b* splice variants are functionally equivalent in nociceptive responses to thermal stimuli in basal conditions. Next, we compared the threshold of paw withdrawal to thermal stimuli among the three genotypes. One paw (ipsilateral) was injected with saline or CFA per mouse, and the contralateral was not subject to injection. We used the Aligned Rank Transform (ART) procedure to assess main and interaction effects of genotype, treatment (saline or CFA), and day post-injection on paw withdrawal thresholds. No significant effect of the interaction among genotype, treatment or day post-injection was found (F_12,252_ = 0.3, p = 0.99). These results suggest that WT, +18a-only, and Δ18a-only mice have similar thresholds to thermal stimuli during CFA-induced inflammation (**Fig. 8A**).

**Figure 8.**
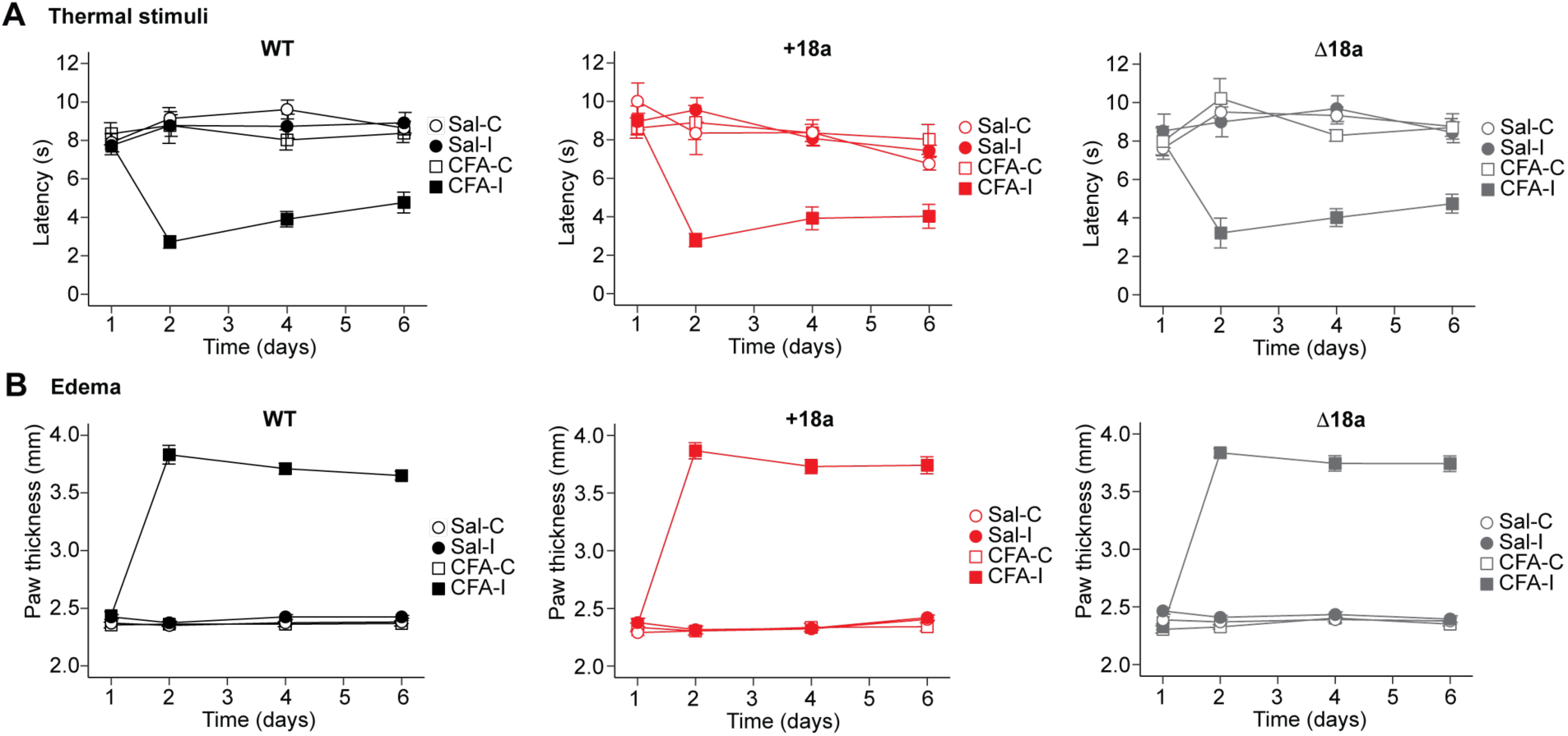
Alternative splicing of exon 18a is indispensable for thermal and edema formation during prolonged inflammation. **A)** Latency of nociceptive responses (paw withdrawal) to thermal stimulation during CFA exposure. After obtaining baseline latencies, left paw was injected with either CFA or saline on day 1. Latency to paw withdrawal was measured every other day for 6 consecutive days on mice of three genotypes. Sal = saline, C = contralateral to the injection site, I = ipsilateral to the injection site.

In addition to measuring thermal hypersensitivity during CFA exposure, we also measured paw thickness as a readout of edema formation. Paw thickness was measured using a digital caliper after performing thermal nociceptive behavior following a similar timeline. In basal conditions, we did not observe a significant difference in paw thickness among the three genotypes (two-way ANOVA, F_2,91_ = 0.250, p = 0.78. **Fig. 8B**). As expected, a significant increase in paw thickness was observed in the presence of CFA compared to saline control for the three genotypes (mixed ANOVA repeated measures, F_1,43_ = 896.8, p < 0.0001). However, no major effect of genotype was observed in the CFA-induced swelling of the paw across the multiple days that we tested (mixed ANOVA repeated measures, F_2,43_ = 0.59, p = 0.28). These results suggest that WT, +18a, and Δ18a mice show similar edema formation in response to CFA injection, further confirming that these splice variants do not play a critical role in CFA-induced inflammation.

Ca_V_2.2 channels are key for capsaicin-induced hypersensitivity to thermal stimuli (2, 3). More recently, Ca_V_2.2 channels expressed in TRPV1-lineage cells underlie most of the thermal hypersensitivity induced by capsaicin (36). TRPV1-lineage cells express both +18a-*Cacna1b* and Δ18a-*Cacna1b* splice variants (37). Therefore, we tested whether these variants are relevant for capsaicin-induced hypersensitivity to thermal stimuli. To test this, we performed the Hargreaves plantar test as described before (36). After baseline measurements of thermal thresholds of mice from the three genotypes (WT, +18a, and Δ18a), one hind paw was injected with either injectable saline or capsaicin. Then, two thermal threshold measurements were obtained at min 15 and 30 after injection. We observed that capsaicin produced a reduction in threshold to thermal stimuli in mice from the three genotypes at 15 min (β = −0.570, SE = 0.174, z = −3.27, p = 0.001, GLMM. **Fig. 9**). However, the mice from the three genotypes showed similar responses to capsaicin (β = 0.159, SE = 0.239, z = 0.665, p = 0.5, GLMM. **Fig. 9**). These results suggest that +18a-*Cacna1b* and Δ18a-*Cacna1*b splice variants contribute equally to capsaicin-induced hypersensitivity to thermal stimulation.

**Figure 9.**
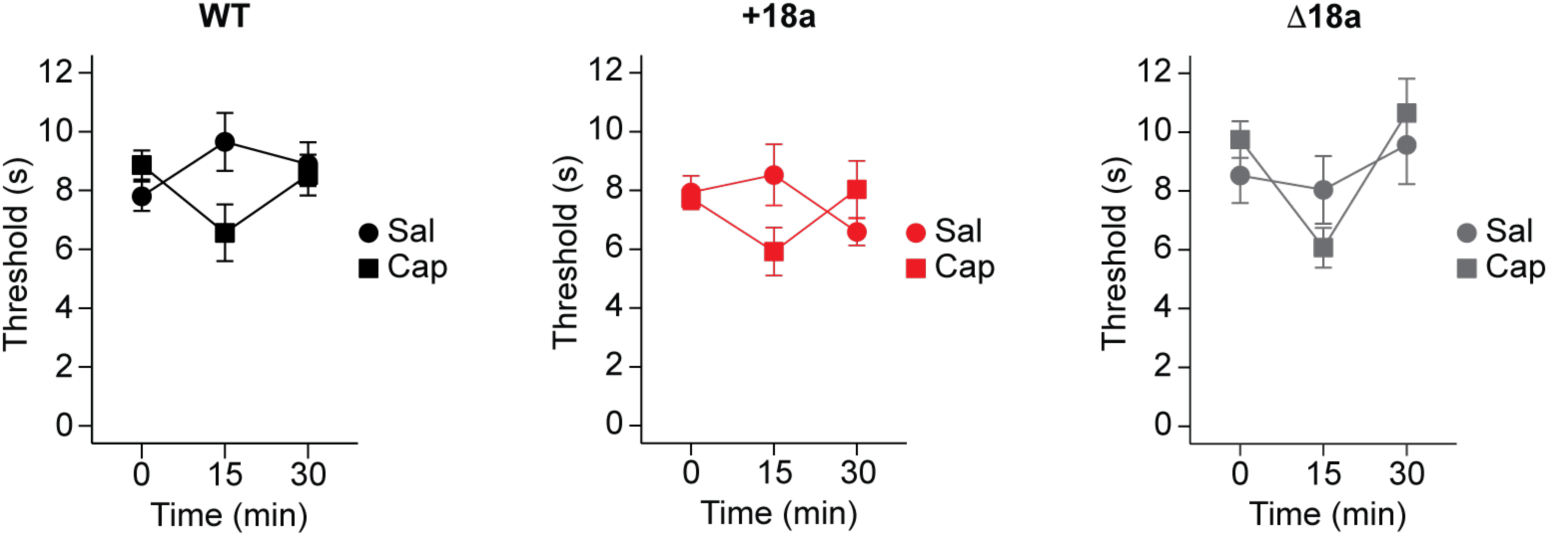
Alternative splicing of exon 18a is indispensable for thermal hypersensitivity during short-term inflammation. Latency to paw withdrawal to thermal stimulation during capsaicin exposure. Immediately after injecting either capsaicin or saline in the left paw, latency to paw withdrawal was measured every 15 min on mice from the three genotypes: WT, *left panel*; +18a, *middle panel*; Δ18a, *right panel*. Data are shown as mean (filled symbols), individual mice (empty symbols), interquartile range (box), median (horizontal line), and upper and lower whiskers, and outliers.

### Alternative splicing of cassette e18a in Cacna1b pre-mRNA is not linked to general locomotion or exploratory behavior

Alternative splicing of the *Cacna1b* pre-mRNA impacts exploratory behavior in basal and under stressful conditions induced by novelty. Specifically, elimination of *Cacan1b* splice variants containing the mutually exclusive exon 37a increases willingness of mice to explore new environments in open spaces and under mild stressful stimuli (18). Furthermore, global elimination of *Cacna1b* results in increased overall locomotion and exploratory behavior. (38) Given this body of evidence, we performed two independent tests, open field maze (OFM) and elevated plus maze (EPM), to assess whether alternative splicing in e18a in the *Cacna1b* pre-mRNA is linked to locomotion and exploratory behavior. In the OFM, we tracked total distance traveled for locomotion and the number of entries and per cent of time spent in the center of the maze versus the outer perimeter for exploratory behavior. No significant differences were found in overall locomotion (distance traveled, mean in m ± SEM: WT = 16.48 ± 0.86, n = 16; e18a = 19.47 ± 1.07, n = 13, Δ18a = 16.7 ± 0.86; F_2,40_ = 3.01, p = 0.06. One-way ANOVA. **Fig. 10A**, *left panel*). No significant differences were observed for exploratory behavior (number of entries into the center, mean ± SEM: WT = 14.4 ± 1.6, n = 16; e18a = 17.5 ± 0.86, n = 13; Δ18a = 17.4 ± 1.7, n = 14; F_2,40_ = 1.44, p = 0.25, One-way ANOVA. **Fig. 10A**, *middle panel*. % of time spent in the center, mean ± SEM, WT = 11.8 ± 2.3, n = 16; e18a = 11.3 ± 1.07, n = 13; Δ18a = 13.1 ± 1.5, n = 14; F_2,40_ = 0.248, p = 0.782, One-way ANOVA. **Fig. 10A**, *right panel*).

**Figure 10.**
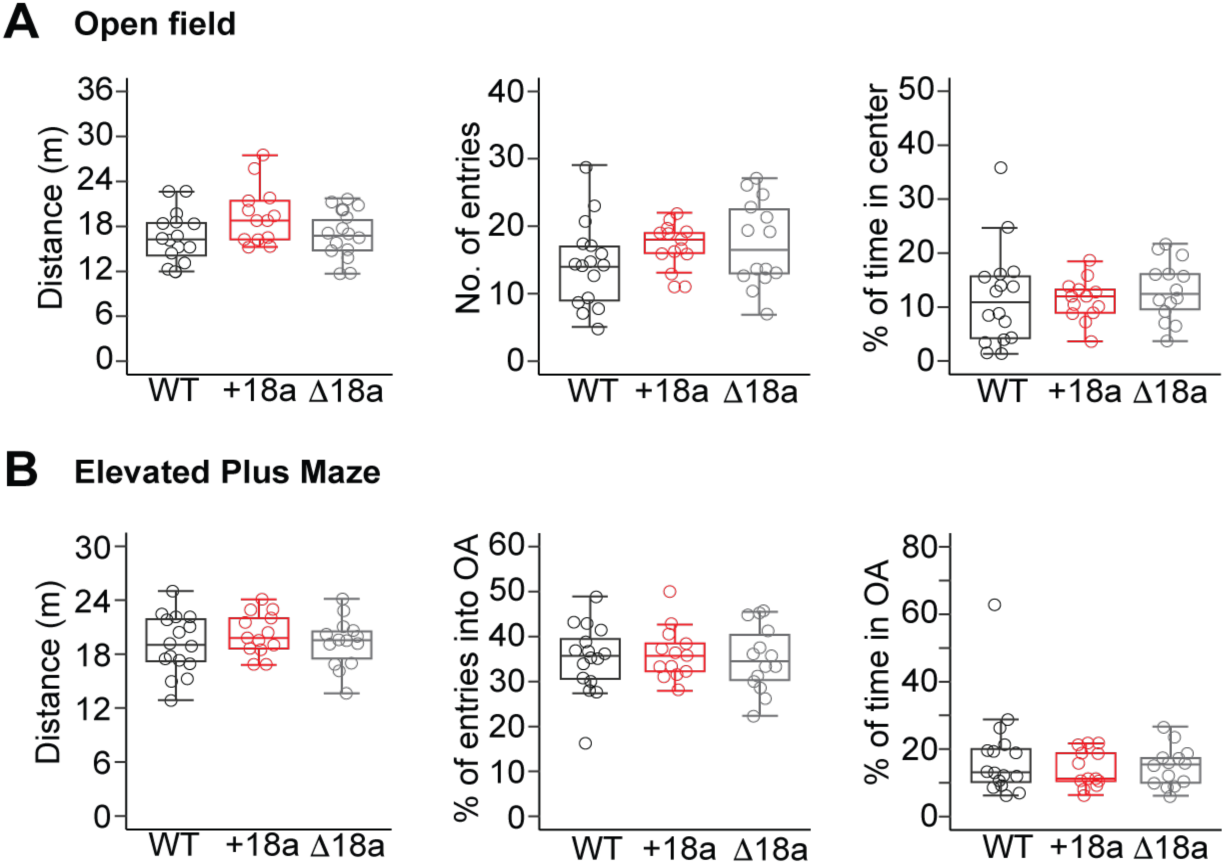
Alternative splicing of exon 18a does not impact overall locomotion and exploratory behavior. **A)** Total distance traveled (*left*), number of entries into the center (*middle*) and % of time in the center (*right*) were measured in mice from the three genotypes. Total distance traveled (left), % of entries into the open arms (OA), and % of time in OA were assessed in mice from the three genotypes.

We also tracked the behavior of mice from the three genotypes in the EPM. We measured distance traveled, number of entries into the open arms, and percentage of time spent in the open arms. Similar to our results in the OFM, no significant differences were found in overall locomotion (distance traveled, mean in m ± SEM: WT = 19.1 ± 0.82, n = 16; e18a = 20.24 ± 0.65, n = 13, Δ18a = 19.28 ± 0.73; F_2,40_ = 0.64, p = 0.53. One-way ANOVA. **Fig. 10B**, *left panel*). No significant differences were found in exploratory behavior in open arms as measured by percent of entries into the open arms and percent of time spent in the open arms (% of entries into open arms ± SEM: WT = 35.1 ± 1.92, n = 16; e18a = 36.2 ± 1.61, n = 13, Δ18a = 35.1 ± 1.95; F2,40 = 0.12, p = 0.89. One-way ANOVA. **Fig. 10B**, *middle panel*. % of time spent in open arms ± SEM: WT = 18.0 ± 3.42, n = 16; e18a = 14.3 ± 1.56, n = 13, Δ18a = 14.7 ± 1.56; χ² = 0.36, df = 2, p = 0.83, Kruskal-Wallis test. **Fig. 10B**, *right panel*). These results further confirmed our findings in the open field test and support that alternative splicing of e18a is not related to locomotion or exploratory behavior.

## DISCUSSION

This body of work represents the first comprehensive behavioral characterization of mice expressing constitutively +18a-Ca_V_2.2 or Δ18a-Ca_V_2.2 splice variants. We showed that manipulating splicing choice for e18a did not alter total *Cacna1b* mRNA, constitutive and alternative splicing of downstream exons in the *Cacna1b* pre-mRNA or total Ca_V_2.2 protein levels. Alternative splicing of e18a selectively impacts aversive associative learning (trace fear conditioning and contextual fear conditioning), but not spatial learning, working memory, nociception in basal conditions and during inflammation, locomotion or exploratory behavior. This points at a specific role of *Cacna1b* splicing at e18a on behavior.

Our results show that +18a-only mice have reduced freezing during trace and contextual phases of trace fear conditioning, indicating impaired temporal association to aversive stimuli and contextual memory. By contrast, Δ18a-only have enhanced freezing during the trace interval, suggesting facilitated encoding of temporal association. These bidirectional effect supports that balanced expression of both splice variants may be required for optimal associative learning. Given that +18a-Ca_V_2.2 is enriched in CCK^+^INs and Δ18a-Ca_V_2.2 predominates in pyramidal neurons (18), these results imply that splicing of cassette exon 18a regulates inhibitory-excitatory interplay during memory acquisition to aversive stimulation.

Consistent with prior studies showing that Ca_V_2.2 channels are relevant for cognitive tasks (13, 14), our studies add a new layer of regulation by linking splicing specificity to learning and memory. The selective phenotype in trace fear conditioning, which is strongly linked to hippocampal-amygdala circuits aligns with a dominant role of Ca_V_2.2 channels on GABA release from CCK^+^INs in these regions (6). Mechanistically, inclusion of e18a reduces cumulative inactivation and increases Ca^2+^ current density (23), which might result in increased release probability at inhibitory synapses. This opens the possibility that e18a splicing modulates GABA release from CCK^+^INs, thereby affecting the timing of network inhibition during learning. Enhanced freezing during trace conditioning in the Δ18a-only mice might reflect stronger or more precisely timed pyramidal neuron excitation due to reduced inhibitory control.

In our studies, we did not observed differences in nociceptive responses to thermal stimuli during basal conditions and in acute (capsaicin) or chronic (CFA) inflammation among mice from the three genotypes. This suggests that e18a splicing does not modulate pain signaling unlike other splicing events in the *Cacna1b* pre-mRNA (e.g. e37a and e37b) (16). Similarly, normal locomotor and exploratory behaviors in the OFM and EPM indicate that altered associative learning in our mouse lines is not secondary to motor defects.

In our studies, we used mouse models with global restriction of splice choice, so the behavioral phenotypes observed in our mouse lines were characterized at the whole-animal level. Thus, cell-type specific electrophysiology (e.g CCK^+^IN-Pyramidal neurons) and behavior are needed to confirm synaptic mechanisms and their relationship to our behavioral outcomes. It also remains to be tested whether stress, hormones, or neuromodulators dynamically regulate e18a inclusion in the *Cacna1b* pre-mRNA. In conclusion, alternative splicing of e18a in the Cacan1b pre-mRNA acts as molecular switch to control associative learning. These findings establish a direct behavioral role *Cacna1b* splice variants in fine-tuning learning and memory expanding our understanding of the role Ca_V_2.2 channels on cognitive functions.

## LIST OF ABBREVIATIONS

*Cacna1b*

Ca_V_2.1

CaV2.2

CaV2.3

+18a

Δ18a

N-type

CCK+INs

Synprint

SNARE

OFM

EPM

BM

TFC

MWM

## DECLARATIONS

### Ethics approval and consent to participate

All of our experiments including animals were approved by the Institutional Animal Care and Use Committee from the University of New Hampshire and Brown University.

### Consent for publication

Not applicable.

### Availability of data and materials

Details of our mousselines are reported in MGI (https://www.informatics.jax.org/) and are available upon request.

## Author’s contributions

SD, CA, designed and performed cognitive and exploratory behavior tasks; BA and BA, performed molecular biology experiments BA; BA, FM, and TV performed nociceptive behavior tasks, CC performed behavior assays; AA designed the study. All authors contributed to data analysis and writing.

## Acknowledgements.

We thank Brianna LaCarubba and Melanie Bertolino for their technical support.

## REFERENCES

1. Dolphin AC, Lee A. Presynaptic calcium channels: specialized control of synaptic neurotransmitter release. Nat Rev Neurosci. 2020;21:213–229.

2. DuBreuil DM, Lopez Soto EJ, Daste S et al. Heat But Not Mechanical Hypersensitivity Depends on Voltage-Gated Ca_V_2.2 Calcium Channel Activity in Peripheral Axon Terminals Innervating Skin. J Neurosci. 2021;41:7546–7560.

3. Salib A-MN, Crane MJ, Lee SH, Wainger BJ, Jamieson AM, Lipscombe D. Interleukin-1α links peripheral Ca_V_2.2 channel activation to rapid adaptive increases in heat sensitivity in skin. bioRxiv. 20242023.12.17.572072.

4. Salib A-MN, Crane MJ, Jamieson AM, Lipscombe D. Peripheral Ca_V_2.2 Channels in the Skin Regulate Prolonged Heat Hypersensitivity during Neuroinflammation. eNeuro. 2024;11:ENEURO.0311-24.2024.

5. Brimblecombe KR, Gracie CJ, Platt NJ, Cragg SJ. Gating of dopamine transmission by calcium and axonal N-, Q-, T- and L-type voltage-gated calcium channels differs between striatal domains. J Physiol. 2015;593:929–946.

6. Blazon M, LaCarubba B, Bunda A et al. N-type calcium channels control GABAergic transmission in brain areas related to fear and anxiety. OBM Neurobiol. 2021;5

7. Armstrong C, Soltesz I. Basket cell dichotomy in microcircuit function. J Physiol. 2012;590:683–694.

8. Bartos M, Elgueta C. Functional characteristics of parvalbumin- and cholecystokinin-expressing basket cells. J Physiol. 2012;590:669–681.

9. Földy C, Neu A, Jones MV, Soltesz I. Presynaptic, activity-dependent modulation of cannabinoid type 1 receptor-mediated inhibition of GABA release. J Neurosci. 2006;26:1465–1469.

10. Hefft S, Jonas P. Asynchronous GABA release generates long-lasting inhibition at a hippocampal interneuron-principal neuron synapse. Nat Neurosci. 2005;8:1319–1328.

11. Lenkey N, Kirizs T, Holderith N et al. Tonic endocannabinoid-mediated modulation of GABA release is independent of the CB1 content of axon terminals. Nat Commun. 2015;6:6557.

12. Szabo Z, Obermair GJ, Cooper CB, Zamponi GW, Flucher BE. Role of the synprint site in presynaptic targeting of the calcium channel CaV2.2 in hippocampal neurons. Eur J Neurosci. 2006;24:709–718.

13. Zhou Y, Niimi K, Li W, Takahashi E. Effects of a Cav2. 2 inhibitor on spatial and non-spatial short-term memory.

14. Jeon D, Kim C, Yang Y-M et al. Impaired long-term memory and long-term potentiation in N-type Ca2+ channel-deficient mice. Genes Brain Behav. 2007;6:375–388.

15. Lipscombe D, Allen SE, Toro CP. Control of neuronal voltage-gated calcium ion channels from RNA to protein. Trends Neurosci. 2013;36:598–609.

16. Andrade A, Denome S, Jiang Y-Q, Marangoudakis S, Lipscombe D. Opioid inhibition of N-type Ca2+ channels and spinal analgesia couple to alternative splicing. Nat Neurosci. 2010;13:1249–1256.

17. Raingo J, Castiglioni AJ, Lipscombe D. Alternative splicing controls G protein-dependent inhibition of N-type calcium channels in nociceptors. Nat Neurosci. 2007;10:285–292.

18. Bunda A, LaCarubba B, Bertolino M et al. Cacna1b alternative splicing impacts excitatory neurotransmission and is linked to behavioral responses to aversive stimuli. Mol Brain. 2019;12:81.

19. Bunda A, Andrade A. BaseScope™ Approach to Visualize Alternative Splice Variants in Tissue. Methods Mol Biol. 2022;2537:185–196.

20. Pan JQ, Lipscombe D. Alternative splicing in the cytoplasmic II-III loop of the N-type Ca channel alpha 1B subunit: functional differences are beta subunit-specific. J Neurosci. 2000;20:4769–4775.

21. Allen SE, Toro CP, Andrade A, López-Soto EJ, Denome S, Lipscombe D. Cell-Specific RNA Binding Protein Rbfox2 Regulates Ca_V_2.2 mRNA Exon Composition and Ca_V_2.2 Current Size. eNeuro. 2017;4:ENEURO.0332-16.2017.

22. Harkins AB, Cahill AL, Powers JF, Tischler AS, Fox AP. Deletion of the synaptic protein interaction site of the N-type (CaV2.2) calcium channel inhibits secretion in mouse pheochromocytoma cells. Proc Natl Acad Sci U S A. 2004;101:15219–15224.

23. Thaler C, Gray AC, Lipscombe D. Cumulative inactivation of N-type CaV2.2 calcium channels modified by alternative splicing. Proc Natl Acad Sci U S A. 2004;101:5675–5679.

24. Malhotra S, Donneger F, Farrell JS, Dudok B, Losonczy A, Soltesz I. Integrating endocannabinoid signaling, CCK interneurons, and hippocampal circuit dynamics in behaving animals. Neuron. 2025;113:1862–1885.

25. Nguyen R, Sivakumaran S, Lambe EK, Kim JC. Ventral hippocampal cholecystokinin interneurons gate contextual reward memory. iScience. 2024;27:108824.

26. Rangel Guerrero DK, Balueva K, Barayeu U et al. Hippocampal cholecystokinin-expressing interneurons regulate temporal coding and contextual learning. Neuron. 2024;112:2045–2061.e10.

27. Adams DJ, Quail MA, Cox T et al. A genome-wide, end-sequenced 129Sv BAC library resource for targeting vector construction. Genomics. 2005;86:753–758.

28. Transparency Is the Key to Quality [editorial]. J Biol Chem 2015;290(50):29692.

29. Ahmed MS, Siegelbaum SA. Recruitment of N-Type Ca(2+) channels during LTP enhances low release efficacy of hippocampal CA1 perforant path synapses. Neuron. 2009;63:372–385.

30. Maximov A, Bezprozvanny I. Synaptic targeting of N-type calcium channels in hippocampal neurons. J Neurosci. 2002;22:6939–6952.

31. Barnes CA. Memory deficits associated with senescence: a neurophysiological and behavioral study in the rat. J Comp Physiol Psychol. 1979;93:74–104.

32. Gandini MA, Zamponi GW. The N-type calcium channel rises from the ashes. J Clin Invest. 2025;135:e189308.

33. Tang C, Gomez K, Chen Y, et al. C2230, a preferential use- and state-dependent CaV2.2 channel blocker, mitigates pain behaviors across multiple pain models. J Clin Invest. 2024;135:e177429.

34. Altier C, Dale CS, Kisilevsky AE et al. Differential role of N-type calcium channel splice isoforms in pain. J Neurosci. 2007;27:6363–6373.

35. Asadi S, Javan M, Ahmadiani A, Sanati MH. Alternative splicing in the synaptic protein interaction site of rat Ca(v)2.2 (alpha (1B)) calcium channels: changes induced by chronic inflammatory pain. J Mol Neurosci. 2009;39:40–48.

36. Meir RY, Sisti MS, Andrade A, Lipscombe D. Trpv1-dependent Cacna1b gene inactivation reveals cell-specific functions of Ca_V_2.2 channels in vivo. Channels (Austin). 2026;20:2594893.

37. Bell TJ, Thaler C, Castiglioni AJ, Helton TD, Lipscombe D. Cell-specific alternative splicing increases calcium channel current density in the pain pathway. Neuron. 2004;41:127–138.

38. Beuckmann CT, Sinton CM, Miyamoto N, Ino M, Yanagisawa M. N-type calcium channel alpha1B subunit (Cav2.2) knock-out mice display hyperactivity and vigilance state differences. J Neurosci. 2003;23:6793–6797.

